# Sex-specific inhibition and stimulation of worker-reproductive transition in a termite

**DOI:** 10.1101/120352

**Authors:** Qian Sun, Kenneth F. Haynes, Jordan D. Hampton, Xuguo Zhou

## Abstract

In social insects, the postembryonic development of individuals exhibits strong phenotypic plasticity in response to environment, thus generating the caste system. Different from eusocial Hymenoptera, in which queens dominate reproduction and inhibit worker fertility, the primary reproductive caste in termites (kings and queens) can be replaced by neotenic reproductives derived from functionally sterile individuals. Feedback regulation of nestmate differentiation into reproductives has been suggested, but the sex-specificity remains inconclusive. In the eastern subterranean termite, *Reticulitermes flavipes*, we tested the hypothesis that neotenic reproductives regulate worker-reproductive transition in a sex-specific manner. With this *R. flavipes* system, we demonstrate a sex-specific regulatory mechanism with both inhibitory and stimulatory functions. Neotenics inhibit workers of the same sex from differentiating into additional reproductives, but stimulate workers of the opposite sex to undergo this transition. Furthermore, this process is not affected by the presence of soldiers. Our results highlight the extraordinary reproductive plasticity of termites in response to social cues, and provide insights into the regulation of reproductive division of labour in a hemimetabolous social insect.

## Introduction

Developmental plasticity plays an important role in the reproductive division of labour in social insects (Page & Amdam 2007). Caste differentiation in eusocial colonies is usually dependent on social stimuli as well as other environmental cues (Hartfelder & Engels 1998; Korb & Hartfelder 2008). Although a fertilized egg is thought to be totipotent and able to develop into any caste, only a few individuals eventually become reproductives. For example, female honeybee larvae that are fed with royal jelly develop into queens, while others become workers (Kucharski et al. 2008). The presence of queens in social Hymenoptera also inhibits worker reproduction by directly supressing their ovarian development, or through policing behaviour (Le Conte & Hefetz 2008).

As with most social insects, termites have caste systems resulting from developmental plasticity. In contrast to social Hymenoptera, hemimetabolous termites have both males and females for all castes. Termite colonies are typically founded by a pair of dispersing alates, which become the primary reproductives, i.e., kings and queens. In many “higher” termite genera (Termitidae) and most “lower” termite genera (all other termite families), workers and nymphs can differentiate into neotenic reproductives (ergatoids and nymphoids, respectively) and reproduce in the natal colony (Myles 1999; Roisin 2000; Roisin & Korb 2011). Neotenic reproduction is implicated to play a critical role in the early evolution of termite eusociality (Myles 1999). The fact that neotenics develop in response to orphaning (the absence of reproductives) has led to the prevailing hypothesis that fertile reproductives would inhibit sexual development (Long, Thorne & Breisch 2003; Matsuura et al. 2010; Moore 1974; Noirot 1990). A few studies, however, proposed the stimulatory effects of reproductive individuals on this process. For example, in *Mastotermes darwiniensis*, neotenic reproductives were produced in the presence rather then the absence of neotenics, and female neotenics exhibited stronger stimulatory activities on workers of both sexes than males (Watson, Metcalf & Sewell 1975). In *Kalotermes flavicollis*, the production of female neotenics was promoted by the presence of a single male neotenic, while the stimulatory effect was not observed from female neotenics (Lüscher 1964). Although kings and queens can both be replaced by neotenics, the sex-specificity for either inhibition or potential stimulation is not conclusive in termites.

*Reticulitermes*, one of the most widely distributed termite genera in the world with substantial economic and ecological importance (Su, Scheffrahn & Cabrera 2001), is an ideal system to study developmental plasticity. *Reticulitermes* workers have three morphologically, behaviourally and functionally distinct developmental trajectories. They can undergo *status quo* moults and remain as workers, differentiate into pre-soldiers followed by an additional moult into soldiers, or develop into neotenic reproductives (i.e., ergatoids) (Lainé & Wright 2003; Zhou, Oi & Scharf 2006). Our preliminary study in the eastern subterranean termite *Reticulitermes flavipes* indicated that worker-reproductive transition was a lengthy process under orphaning condition (30-90 days). If one of the reproductives (e.g., queen) is lost, a stimulatory function from the remaining reproductive (e.g., king) that promotes the formation of neotenic reproductives of the missing sex (e.g., female ergatoid) would be beneficial to the colony. We hypothesized that worker-reproductive transition is regulated in a sex-specific manner in *R. flavipes*. Specifically, reproductive individuals inhibit same-sex workers, but stimulate opposite-sex workers to differentiate into ergatoids. To test this, we evaluated ergatoid formation in response to the presence or absence of male or female reproductives. As soldier caste is present in all termite species (Tian & Zhou 2014), and previous studies suggest that soldiers potentially promote differentiation of reproductives (Watanabe et al. 2014), we also examined the effect of soldiers on ergatoid formation.

## Methods

### Study System

Colonies of *R. flavipes* were collected from the Arboretum (Lexington, Kentucky, USA), and the Red River Gorge area, Daniel Boone National Forest (Slade, Kentucky, USA). Colonies consisted of workers, soldiers and nymphs at the time of collection. Freshly collected termites were kept in Petri dishes and fed on moistened unbleached paper towel for one to two weeks. Neotenic-headed colonies were obtained by transferring field collected termites to closed plastic boxes (45.7 × 30.6 × 15.2 cm) filled with moistened wood mulch and pinewood blocks and maintained for 6 months, when eggs and larvae appeared indicating the presence of fertile neotenic reproductives. Primary-headed colonies were established by pairing female and male sibling alates collected in Lexington, Kentucky, and they had been maintained in laboratory for 5 years by the time experiments started. All stock colonies and experimental termites were maintained at 27 ± 1°C in complete darkness.

### Bioassay to Test Sex-specific Regulation

Fertile ergatoids of different sexes were used to test their influences on worker-reproductive transition. These ergatoids were obtained from an orphaning assay, in which groups of 100 workers were kept in Petri dishes (6.0 cm in diameter, 1.5 cm in height) lined with moistened unbleached paper towel for 60 days. Ergatoids that were actively reproducing (i.e., with eggs present in dishes) were used in the subsequent assay.

The same set-up was used to test how workers differentiate in response to ergatoids (Fig. 1). Groups of 100 workers were kept with: 1) no reproductives (F-M-); 2) one female ergatoid (F+M-); 3) one male ergatoid (F-M+); or 4) one pair of ergatoids (F+M+). The ergatoids and workers were from the same colony in each group. Fourteen replications using three colonies were conducted, with one colony originally primary-headed and two colonies originally neotenic-headed. Worker differentiation was observed daily for 60 consecutive days, and newly formed ergatoids were removed and their sex was determined. Ergatoids were recognized by slightly heavier cuticle pigmentation, elongated abdomen, and wider thorax than workers (Fig. 2). Female ergatoids were distinguished from males by their enlarged 7th abdominal sternite. We also removed eggs, newly formed pre-soldiers and any inter-caste (individuals with the degree of mandible development intermediate between workers and pre-soldiers) every day to prevent their potential influence on worker development.

**Figure 1.**
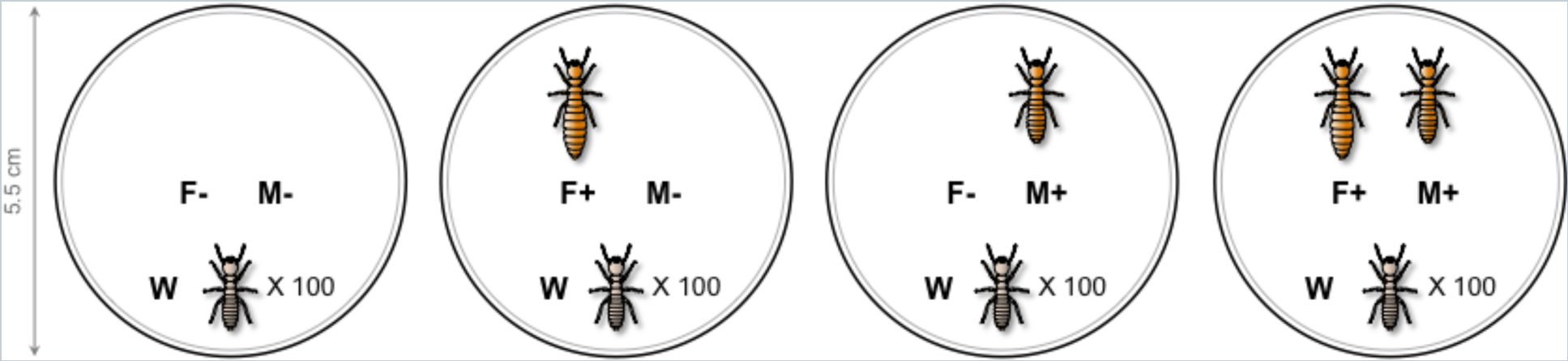
Experimental set-up. Each group of 100 workers were placed in a Petri dish lined with moistened unbleached paper. Workers were kept with no reproductives (F-M-), one female ergatoid (F+M-), one male ergatoid (F-M+), or one pair of ergatoids (F+M+). Newly formed ergatoids, pre-soldiers, and eggs laid by reproductives were removed every day for 60 consecutive days. FR: female ergatoid reproductive; MR: male ergatoid reproductive; W: worker. Ergatoid reproductives used in the assay were actively reproducing, and female ergatoids were physogastric.

**Figure 2.**
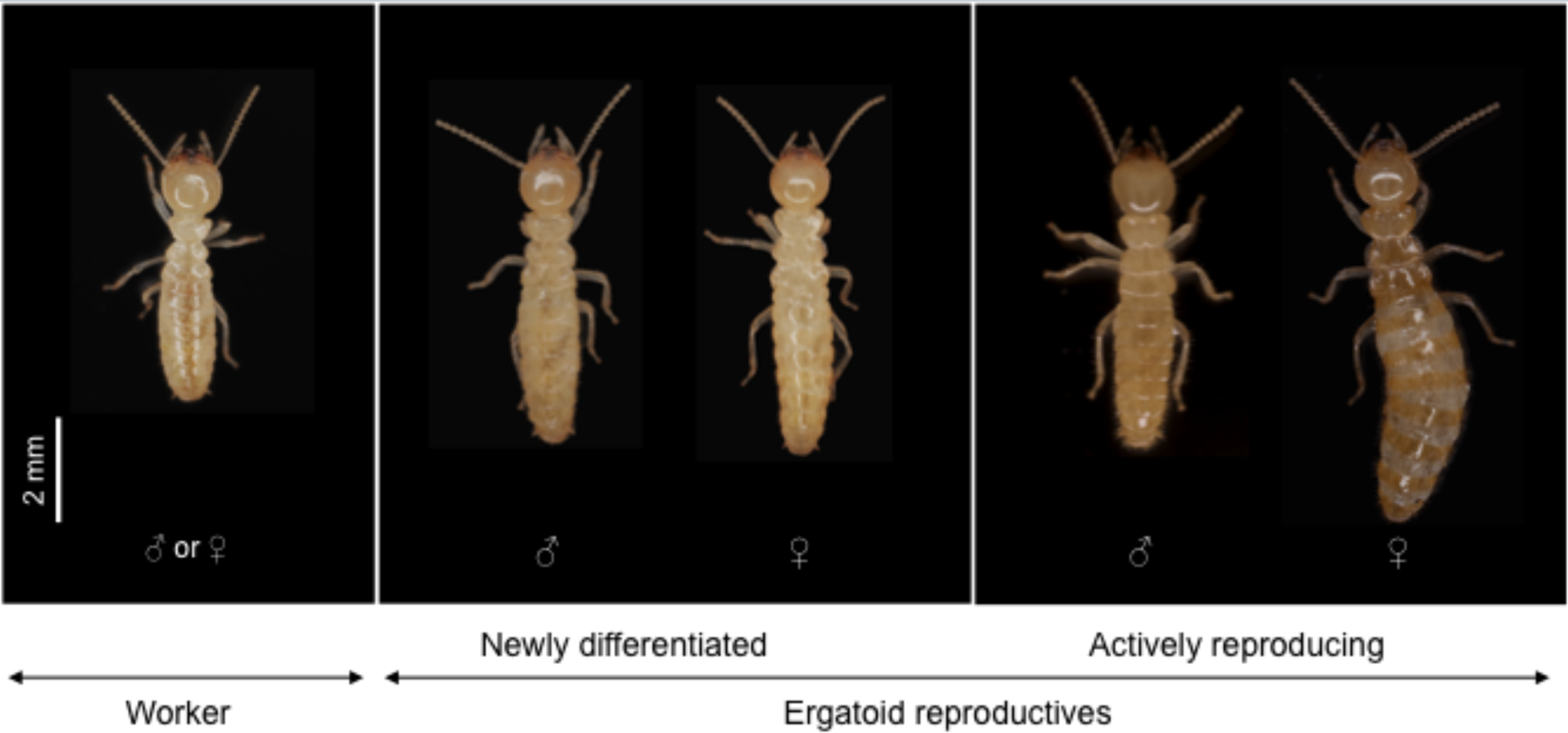
Photographs of a worker, young ergatoids, and mature ergatoids. The young ergatoids were about 10 days post differentiation. The mature ergatoids were six months post differentiation and actively reproducing.

### Bioassays to Test Soldier Effect

Two orphaning assays were conducted to test influence of soldier caste on ergatoid differentiation. The first assay (“single-soldier orphaning assay”) simulated natural orphaning condition where freshly collected workers were separated into groups of 100 individuals, and each group was placed with one soldier (Soldier+) or no soldier (Soldier-). Termites were maintained in Petri dishes (6.0 cm in diameter, 1.5 cm in height) lined with moistened unbleached paper towel as food source. Termites were allowed to undergo changes in caste differentiation in the dishes without disturbance, and caste composition and mortality of each group was documented at the end of 60 days. A total of 42 and 45 replicates from four colonies were conducted for soldier+ and soldier-treatment, respectively.

We further performed the second assay (“multiple-soldier orphaning assay”), which was conducted with an increased soldier stimulus and observed daily for 60 consecutive days. Groups of 100 workers were isolated from neotenic-headed colonies, and they were companioned with either four soldiers (Soldier++) or no soldier (Soldier-). Newly differentiated individuals (ergatoids and pre-soldiers) were removed to prevent their potential influence on worker development, and an equal number of workers to the removed individuals were added to the group to keep group size consistent. Termites were maintained in Petri dishes (3.5 cm in diameter, 1.5 cm in height) provided with moistened unbleached paper towel. A total of 10 replicates from two colonies were conducted for both soldier++ and soldier-treatments.

### Data Analyses

Data were analysed using Statistix 10 (Analytical Software, Tallahassee, FL, USA). In the assay that tested sex-specific regulation, Wilcoxon rank-sum test was performed on the cumulative numbers ergatoids between each treatment and the control on every assay day. In both single- and multiple-soldier orphaning assays, unpaired *t*-test was performed on numbers of differentiated individuals and mortality. To obtain values that fit the assumptions of parametric test, data were transformed through square root (x + 0.5) on the combined numbers of pre-soldiers and soldiers in the single-soldier orphaning assay, and numbers of female and male ergatoids in the multiple-soldier orphaning assay. Because the pattern of regulation was consistent in all colonies, results were pooled for statistical analyses.

## Results

### Ergatoid Formation in Response to Reproductives of Different sexes

Under orphaning condition (F-M-), 4.1 ± 1.3% and 2.5 ± 0.7% of workers differentiated into female and male ergatoids, respectively, in 60 days (mean ± SEM, Fig. 3). The presence of a single female (F+M-) or a pair of ergatoids (F+M+) significantly inhibited the formation of additional female ergatoids (0.2 ± 0.2% and 0.1 ± 0.1%, respectively); however, the presence of a single male ergatoid (F-M+) significantly stimulated the formation of female ergatoid (26.8 ± 3.0%) (Fig. 3(a); *P* < 0.01, Wilcoxon rank-sum test, one-sided, n = 14). Similarly, significantly fewer male ergatoids differentiated in the presence of a single male (F-M+) or a pair of ergatoids (F+M+) (0% and 0.1 ± 0.1%, respectively), while significantly more of them formed when a single female ergatoid was present (F+M-) (14.3 ± 1.6%) (Fig. 3(b); *P* < 0.01, Wilcoxon rank-sum test, one-sided, n = 14).

**Figure 3.**
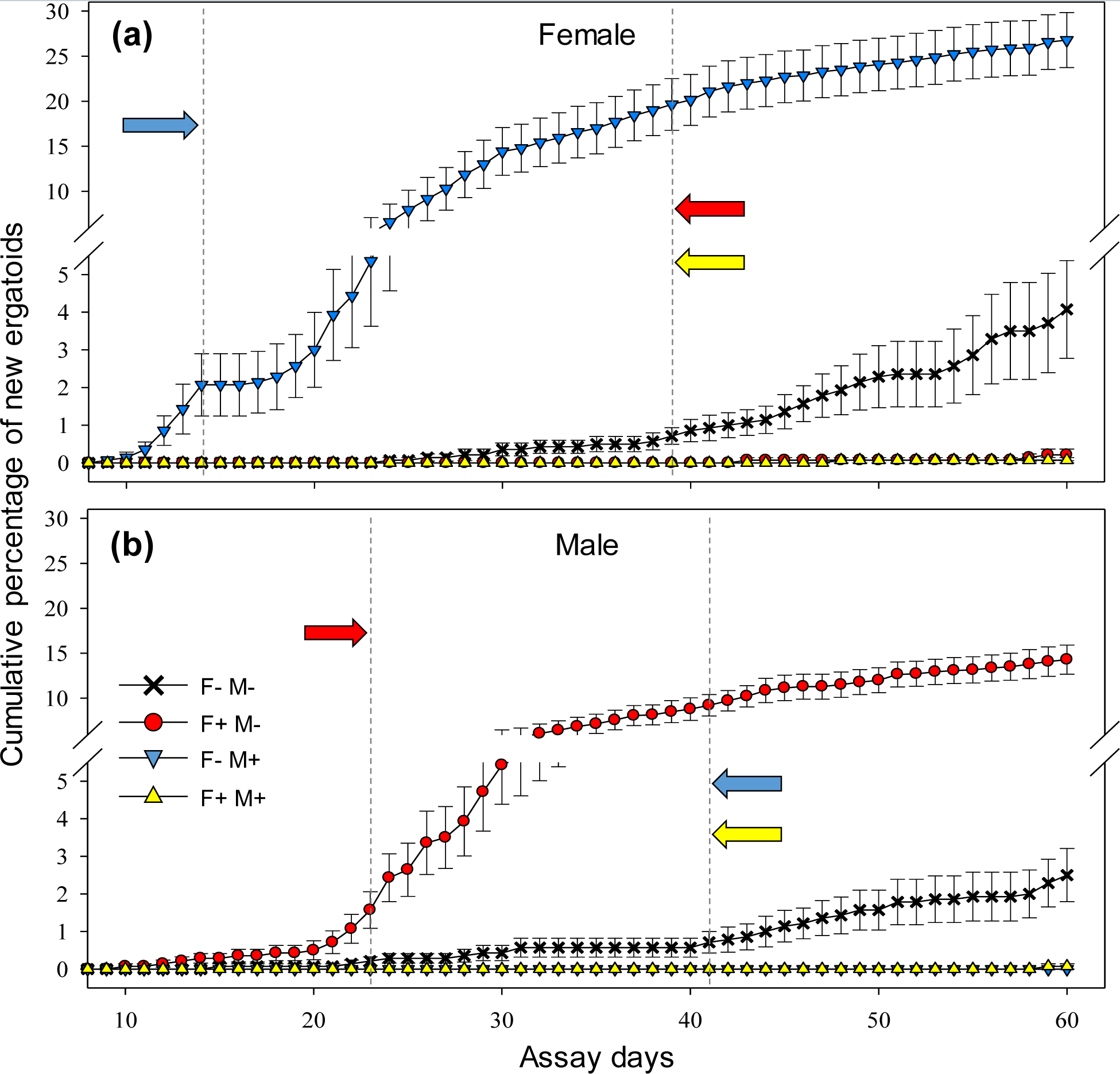
Ergatoid formation in response to fertile ergatoids of different sexes. Cumulative percentage of newly differentiated female (a) and male (b) ergatoids is shown (mean ± SEM). Stimulation (forward arrows) and inhibition (reverse arrows) refer to significantly more and fewer ergatoids formed, respectively, relative to control (F-M-), and dash line next to the tip of arrow indicates the day when the significant difference started (Wilcoxon rank-sum test, one-sided, *P* < 0.01; n = 14 for all treatments). Symbols and arrows of the same colour correspond to the same treatment.

Within 60 days, female and male ergatoids differentiated in 10 and 9 replicates, respectively, out of 14 total replicates under the orphaning condition (F-M-). The formation of ergatoids in all 14 replicates was stimulated by a single ergatoid of the opposite sex. Under this stimulation, developmental time for the first ergatoid was significantly reduced (female: 19 ± 3.3 days, n = 14 in F-M+, compared with 38 ± 3.7 days, n = 10 in F-M-; male: 21 ± 1.8 days, n = 14 in F+M-, compared with 35 ± 5.4 days, n = 9 in F-M-; Wilcoxon rank-sum test, one-sided, *P* < 0.01). There was no significant difference on mortality within 60-day assay period among treatments (Fig. 4; ANOVA, *F_3,52_* = 1.5, *P* > 0.05; n = 14).

**Figure 4.**
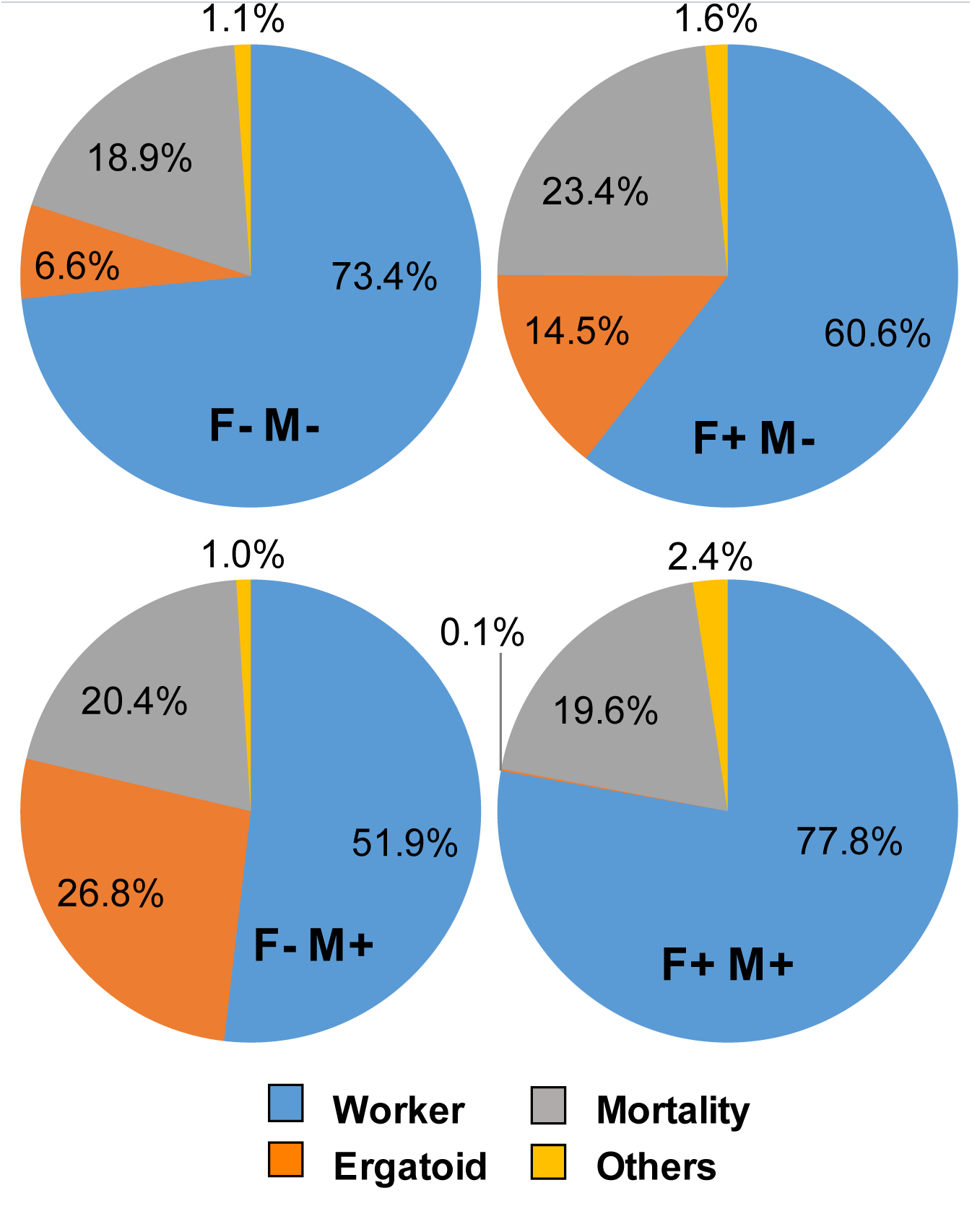
Developmental endpoint of workers in 60 days. “Others” includes pre-soldiers and inter-castes, and the latter refers to individuals with the degree of mandible development between workers and pre-soldiers after moulting. There was no significant effect of treatment on mortality (ANOVA, *F*_*3,52*_ = 1.5, *P* > 0.05; n = 14 pooled from three colonies).

### Ergatoid Formation in Response to Soldier Caste

In both assays, ergatoids were differentiated from workers in response to orphaning condition, and there were no significant effects of soldiers on ergatoid formation (Fig. 5). In the single-soldier orphaning assay, soldier caste did not significantly influence the number of female or male ergatoids in 60 days (Fig. 5(a); female: t_85_ = 1.64, *P* > 0.05; male: t_85_ = 1.64, *P* > 0.05; unpaired *t*-test, two-sided, n = 42 for Soldier+; n = 45 for Soldier-). At the end of the assay, mortality between Soldier+ and Soldier-groups were not significantly different (Fig. 5(b); t_42_ = 0.67, *P* > 0.05; unpaired *t*-test, two-sided, n = 22 randomly selected from both soldier+ and soldier-groups). The presence of one soldier significantly supressed the differentiation of pre-soldiers and soldiers, and the increased total number of pre-soldiers and soldiers are 0.43 ± 0.10 in Soldier+ groups and 0.91 ± 0.14 in Soldier-groups (t_85_ = 2.71, *P* < 0.01; unpaired *t*-test, two-sided; data were transformed (square root (x + 0.5)); n = 42 for Soldier+, n = 45 for Soldier-). Similarly, in the multiple-soldier orphaning assay, the soldier caste did not significantly influence the accumulative number of female or male ergatoids in 60 days (Fig. 5(c); female: t_18_ = 0.57, *P* > 0.05; male: t_18_ = 0.44, *P* > 0.05; unpaired *t*-test, two-sided; data were transformed (square root (x +0.5)); n = 10 for both Soldier++ and Soldier-groups). Mortality was not significantly influenced by the presence of 4 soldiers (Fig. 5(d); t_18_ = 1.02, *P* > 0.05; unpaired *t*-test, two-sided, n = 10 for both Soldier++ and Soldier-groups).

**Figure 5.**
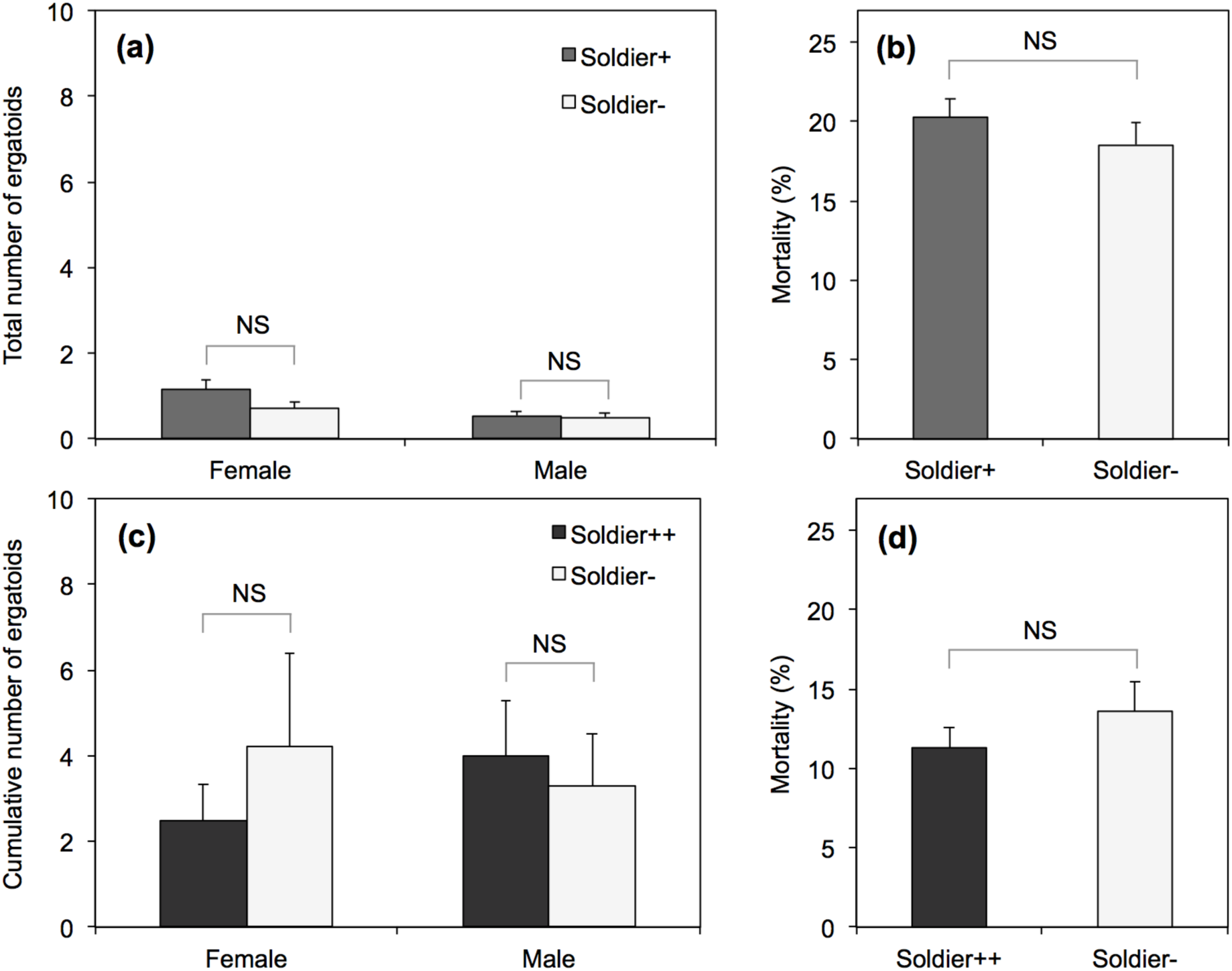
Soldier impacts on ergatoid formation. In a single-solider orphaning assay, total numbers of female and male ergatoids presented (mean ± SEM) (a) and mortality (%, mean ± SEM) (b) between soldier present and absent groups in 60 days are shown. In a multiple-soldier orphaning assay, cumulative numbers of female and male ergatoids differentiated (mean ± SEM) (c) and mortality (%, mean ± SEM) (d) between soldier present and absent groups in 60 days are shown. In the single-soldier assay, newly differentiated ergatoids were left in groups; Soldier+: each group consisted of 100 workers and one soldier at the start of assay; Soldier-: each group consisted of 100 workers only; NS: no significant difference (unpaired *t*-test, two-sided, *P* > 0.05; n = 42 for Soldier+, n = 45 for Soldier-). In the multiple-soldier assay, newly differentiated ergatoids were removed and replaced with workers every day; Soldier++: each group consisted of 100 workers and four soldiers at the start of assay; Soldier-: each group consisted of 100 workers only; NS: no significant difference (unpaired *t*-test, two-sided, *P* > 0.05; n = 10 for both Soldier++ and Soldier-).

## Discussion

These results support our hypothesis that regulation of worker-reproductive transition by fertile reproductives is sex-specific. More importantly, our empirical evidence demonstrated that the dual regulation (inhibition and stimulation) is employed by both sexes. Ergatoid differentiation occurs after more than 30 days in response to orphaning, but can be significantly accelerated by the presence of a potential mate. Such stimulation by the opposite sex benefits the colony by enabling it to resume reproduction soon after the loss of a queen or a king. Inhibition of development by the same sex, on the other hand, prevents unnecessary investments in reproduction, which, in turn, would be a loss in the labour force. This sex-specific regulation suggests that the development of reproductives is strictly dependent on the reproductive needs of the colony.

Neotenic reproduction is common in termites; however, regulation of neotenic differentiation varies among species (Grassé & Noirot 1960; Lüscher 1964; Matsuura et al., 2010; Miyaguni, Sugio & Tsuji 2013; Watson, Metcalf & Sewell 1975). In *R. speratus*, female reproductives inhibit the differentiation of female neotenics, but do not influence the formation of male neotenics (Matsuura et al. 2010). Compared with *R. flavipes*, female ergatoid formation is faster in *R. speratus* in response to orphaning, and the formation of nymphoids are faster than ergatoids in *R. speratus* (Matsuura et al. 2010; Miyata, Furuichi & Kitade 2004). Orphaning assays have also been conducted in other congeneric species including *R. grassei* (Pichon et al. 2007) and *R. urbis* (Ghesini & Marini 2009), which confirmed the inhibitory effect of reproductive pairs, but the regulation by each sex remains unclear. Stimulation by neotenic reproductives has been suggested previously in primitive termite species. In *K. flavicollis*, the formation of female neotenics was stimulated by the extracts of male reproductives (Lüscher 1964). In *M. darwiniensis*, formation of neotenics was promoted in the presence rather than the absence of other neotenics. Although sex-specificity was not confirmed, female neotenics exhibited stronger stimulatory effects than males and the pair (Watson & Abbey 1985; Watson, Metcalf & Sewell 1975).

In comparison, *R. flavipes* neotenics exhibit sex-specific inhibition and stimulation in both sexes (Fig. 6a). Such a regulatory pattern consists of all possible directions of social regulation, therefore presents a model for understanding the pheromonal and developmental mechanisms underlying neotenic reproduction. Given that previous studies on the differentiation of neotenic reproductives were incomplete or inconclusive (Fig. 6b-6d), this study also provides an opportunity for us to re-examine the sex-specificity hypothesis across termite taxa. Foraging populations of *R. flavipes* contain about 2% or less soldiers (Haverty & Howard 1981; Howard & Haverty 1980), while higher proportions (close to 4%) were observed in nest areas where neotenics are present (Howard & Haverty 1980). Soldiers were considered to induce the differentiation of reproductives (Tian & Zhou 2014; Watanabe et al. 2014), however, our results indicated that soldier caste does not play a significant role in regulating ergatoid formation in *R. flavipes*. It is worth noting that if ergatoids were not removed from the groups (single-soldier orphaning assay), the total number of ergatoids was lower than the cumulative number of ergatoids if they were constantly removed (multiple-soldier orphaning assay) within the same period of time. Although the two assays were conducted separately and not entirely comparable, this result could be explained by newly formed ergatoids suppressing formation of additional ergatoids through pheromones or policing behaviour.

**Figure 6.**
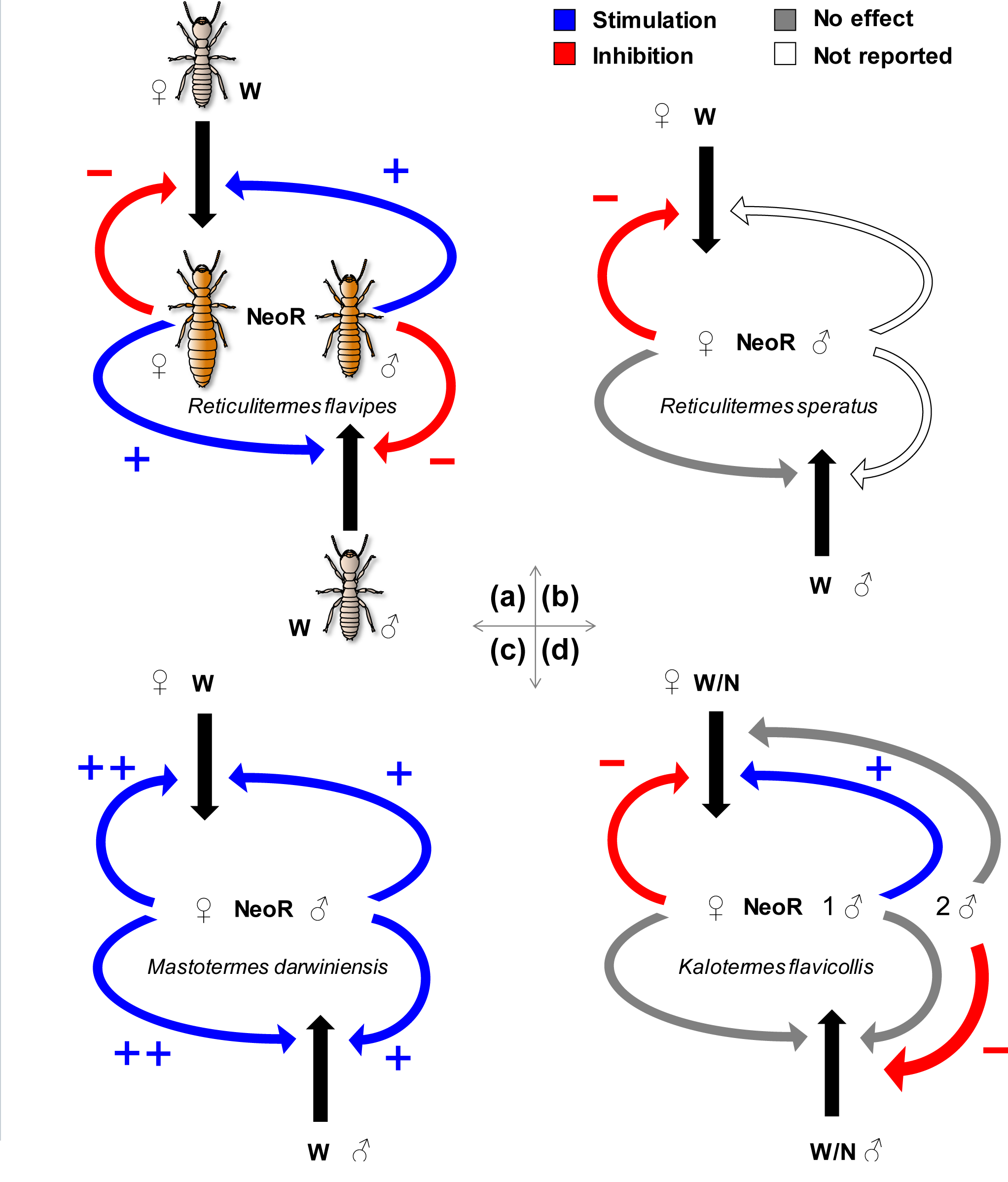
Comparison of feedback regulation of neotenic differentiation in four termites. (a) Sex-specific inhibition and stimulation are demonstrated for both females and males in *R. flavipes* (this study). (b) In *R. speratus*, female neotenics inhibit differentiation of females, but does not influence males; the effects of male neotenics were not reported. (c) In *M. darwiniensis*, both female and male neotenics stimulate neotenic differentiation, but not in a sex-specific manner; females exhibit stronger stimulation. (d) In *K. flavicollis*, female neotenics inhibit differentiation of females, but the effect of males depends on number. One male neotenic shows opposite-sex stimulation, while two males exhibit same-sex inhibition. W: worker; N: nymph; NeoR: neotenic reproductive.

Our study represents a first step in understanding sex-specific worker-reproductive differentiation in response to social cues. The results from this study add a new dimension to the prevailing view that reproductives inhibit worker-reproductive transition in termites (Noirot 1990). Much remains to be investigated about the regulatory mechanisms of caste differentiation, including the identification of inhibitory and stimulatory pheromones from reproductives. The active substances or blends must be sex-specific. The search of reproductive pheromones in termites should include volatile compounds (Matsuura et al. 2010), cuticular hydrocarbons (Liebig, Eliyahu & Brent 2009) as often observed in Hymenoptera (Van Oystaeyen et al. 2014), and proteinaceous secretions (Hanus et al. 2010). The sex-specificity and the dual effect of reproductive cues reflect unique adaptation and regulation of caste differentiation in hemimetabolous termites.

## Acknowledgments

We thank Dr. Li Tian (Pennsylvania State University) for his help with photography, and members of the Zhou lab for their comments and discussion. This study was supported by William L. and Ruth D. Nutting Student Research Grant from the International Union for the Study of Social Insects (North American Section), Kentucky Opportunity Fellowship from the University of Kentucky to Q.S., and the USDA National Institute of Food and Agriculture Hatch project (Accession Number: 1004654) to X.Z. Any opinions, findings, conclusions, or recommendations expressed in this publication are those of the author(s) and do not necessarily reflect the view of the National Institute of Food and Agriculture (NIFA) or the United States Department of Agriculture (USDA). This is publication No. 17-08-002 of the Kentucky Agricultural Experiment Station and is published with the approval of the Director. The granting agencies have no role in the study design, data collection and analysis, decision to publish, or preparation of the manuscript.

## Author Contributions

Q.S., K.F.H. and X.Z. designed the experiments, Q.S. and J.D.H conducted the experiments, Q.S. and K.F.H analysed the data, Q.S. drafted the manuscript, and K.F.H and X.Z. revised the manuscript. All authors approved the final manuscript.

## References

Ghesini, S. & Marini, M. (2009) Caste differentiation and growth of laboratory colonies of *Reticulitermes urbis* (Isoptera, Rhinotermitidae). Insectes Sociaux, 56, 309–318. doi: 10.1007/s00040-009-0025-1

Grassé, P. P. & Noirot, C. (1960) Role respectif des males et des femelles dans la formation des sexués néoténiques chez *Calotermes flavicollis*. Insectes Sociaux, 7,109–123. doi: 10.1007/BF02224075

Hanus, R., Vrkoslav, V., Hrdý, I., Cvačka, J. & Šobotník, J. (2010) Beyond cuticular hydrocarbons: evidence of proteinaceous secretion specific to termite kings and queens. Proceedings of the Royal Society of London B: Biological Sciences, 277, 995–1002. doi: 10.1098/rspb.2009.1857

Hartfelder, K. & Engels, W. (1998) Social insect polymorphism: hormonal regulation of plasticity in development and reproduction in the honeybee. Current Topics in Developmental Biology (eds R.A. Pedersen & G.P. Schatten), pp. 45–78. Academic Press, San Diego.

Haverty, M. & Howard, R. (1981) Production of soldiers and maintenance of soldier proportions by laboratory experimental groups of *Reticulitermes flavipes* (Kollar) and *Reticulitermes virginicus* (Banks) (Isoptera: Rhinotermitidae). Insectes Sociaux, 28, 32–39. doi: 10.1007/BF02223620

Howard, R.W. & Haverty, M.I. (1980) Reproductives in mature colonies of *Reticulitermes flavipes:* abundance, sex-ratio, and association with soldiers. Environmental Entomology, 9, 458–460. doi: http://dx.doi.org/10.1093/ee/9.4.458

Korb, J. & Hartfelder, K. (2008) Life history and development - a framework for understanding developmental plasticity in lower termites. Biological Reviews, 83, 295–313. doi: 10.1111/j.1469-185X.2008.00044.x

Kucharski, R., Maleszka, J., Foret, S. & Maleszka, R. (2008) Nutritional control of reproductive status in honeybees via DNA methylation. Science, 319, 1827–1830. doi: 10.1126/science.1153069

Lainé, L.V. & Wright, D.J. 2003. The life cycle of *Reticulitermes* spp. (Isoptera: Rhinotermitidae): what do we know? Bulletin of Entomological Research, 93, 267–278. doi: https://doi.org/10.1079/BER2003238

Le Conte, Y. & Hefetz. A. (2008) Primer pheromones in social Hymenoptera. Annual Review of Entomology, 53, 523–542. doi: 10.1146/annurev.ento.52.110405.091434

Liebig, J., Eliyahu, D. & Brent, C. (2009) Cuticular hydrocarbon profiles indicate reproductive status in the termite *Zootermopsis nevadensis*. Behavioral Ecology and Sociobiology, 63, 1799–1807. doi: 10.1007/s00265-009-0807-5

Long, C.E., Thorne, B.L. & Breisch N.L. (2003) Termite colony ontogeny: a long-term assessment of reproductive lifespan, caste ratios and colony size in *Reticulitermes flavipes* (Isoptera: Rhinotermitidae). Bulletin of Entomological Research, 93, 439–445. doi: https://doi.org/10.1079/BER2003258

Lüscher, M. (1964). Die spezifische Wirkung männlicher und weiblicher Ersatzgeschlechtstiere auf die Entstehung von Ersatzgeschlechtstieren bei der Termite *Kalotermes flavicollis* (Fabr.) Insectes Sociaux, 11, 79–90. doi: 10.1007/BF02222973

Matsuura, K., Himuro, C., Yokoi, T., Yamamoto, Y., Vargo, E.L. & Keller L. 2010. Identification of a pheromone regulating caste differentiation in termites. Proceedings of the National Academy of Sciences of the United States of America, 107, 12963–12968. doi: 10.1073/pnas.1004675107

Miyaguni, Y., Sugio K. & Tsuji K. (2013) The unusual neotenic system of the Asian dry wood termite, *Neotermes koshunensis* (Isoptera: Kalotermitidae). Sociobiology, 60, 65–68. doi: http://dx.doi.org/10.13102/sociobiology.v60i1.65-68

Miyata, H., Furuichi H. & Kitade O. (2004) Patterns of neotenic differentiation in a subterranean termite, *Reticulitermes speratus* (Isoptera: Rhinotermitidae). Entomological Science, 7, 309–314. doi: 10.1111/j.1479-8298.2004.00078.x

Moore, B. (1974) Pheromones in the termite societies. Pheromones (ed M. Birch), pp. 250–266. North-Holland Publishing, Amsterdam.

Myles, T.G. (1999) Review of secondary reproduction in termites (Insecta: Isoptera) with comments on its role in termite ecology and social evolution. Sociobiology, 33, 1–43.

Noirot, C. 1990. Sexual castes and reproductive strategies in termites. Social Insects (ed W. Engels), pp. 5–35. Springer, Berlin Heidelberg.

Page, R. E. & Amdam G. V. (2007) The making of a social insect: developmental architectures of social design. BioEssays, 29, 334–343. doi: 10.1002/bies.20549

Pichon, A., Kutnik M., Leniaud L., Darrouzet, E., Chaline, N., Dupont S. & Bagnères A. (2007) Development of experimentally orphaned termite worker colonies of two *Reticulitermes* species (Isoptera: Rhinotermitidae). Sociobiology, 50, 1015–1034.

Roisin, Y. (2000) Diversity and evolution of caste patterns. Termites: Evolution, Sociality, Symbioses, Ecology. (eds T. Abe, D.E. Bignell & M. Higashi), pp. 95–119. Springer, Netherlands.

Roisin, Y. & Korb J. (2011) Social organisation and the status of workers in termites. Biology of Termites: a Modern Synthesis. (eds D.E. Bignell, Y. Roisin & N. Lo), pp. 133–164. Springer, Netherlands.

Su, N.-Y., Scheffrahn, R.H. & Cabrera B.J. (2001) Native subterranean termites: *Reticulitermes flavipes* (Kollar), *Reticulitermes virginicus* (Banks), *Reticulitermes hageni* Banks (Insecta: Isoptera: Rhinotermitidae). University of Florida Cooperative Extension Service, Institute of Food and Agricultural Sciences, EDIS,

Tian, L. & X. Zhou. (2014) The soldiers in societies: defense, regulation, and evolution. International Journal of Biological Sciences, 10, 296–308. doi: 10.7150/ijbs.6847

Van Oystaeyen, A., Oliveira, R.C., Holman, L., van Zweden, J.S., Romero, C., et al. (2014) Conserved class of queen pheromones stops social insect workers from reproducing. Science, 343, 287–290. doi: 10.1126/science.1244899

Watanabe, D., Gotoh, H., Miura, T. & Maekawa, K. (2014) Social interactions affecting caste development through physiological actions in termites. Frontiers in Physiology, 5, 127. doi: https://doi.org/10.3389/fphys.2014.00127

Watson, J. & Abbey H.M. (1985) Development of neotenics in *Mastotermes darwiniensis* Froggatt: an alternative strategy. Caste Differentiation in Social Insects (eds J.A.L. Watson, B.M. Okot-Kotber, & C. Noirot), pp. 107–124. Pergamon Press, Oxford.

Watson, J.A.L., Metcalf E.C., & Sewell J.J. (1975) Preliminary studies on the control of neotenic formation in *Mastotermes Darwiniensis* Froggatt (Isoptera). Insectes Sociaux, 22, 415–426. doi: 10.1007/BF02224116

Zhou, X., Oi F.M. & Scharf M.E. (2006) Social exploitation of hexamerin: RNAi reveals a major caste-regulatory factor in termites. Proceedings of the National Academy of Sciences of the United States of America, 103, 4499–4504. doi: 10.1073/pnas.0508866103

